# LagCI Enables Inference of Temporal Causal Relationships from Dense Multi-Omic Time Series

**DOI:** 10.64898/2026.04.15.718654

**Authors:** Yifei Ge, Shunpeng Bai, Zirui Qiang, Yijiang Liu, Yitong Wu, Xiaotao Shen

## Abstract

Inferring causal relationships from time-series data is critical for uncovering the dynamics of biological regulation. However, in multi-omics studies, this task is often hampered by sparse temporal sampling and the limitations of existing methods. To address this, we developed Lagged-Correlation Based Causal Inference (lagCI), a computational framework designed to identify time-lagged associations by combining comprehensive lag-correlation profiling with a robust statistical filtering scheme. Rather than relying on simple cross-correlation, lagCI analyzes the entire correlation profile and applies a quality-scoring system to filter out spurious associations that often plague high-dimensional datasets.

We first tested lagCI on wearable physiological data, where it successfully captured the well-known causal link between physical activity and heart rate, even accounting for variations in lag times between individuals. Moving to high-frequency human multi-omics, we used lagCI to build a directed network of 1,624 molecules connected by over 157,000 predicted interactions. This network didn’t just mirror established biology (such as cytokine-hormone crosstalk); it also pointed to specific molecular hubs that seem to orchestrate the timing of metabolic and immune responses.

Overall, lagCI provides a data-driven way to extract temporal insights from dense longitudinal omics. We’ve made the tool available as an R package with multiple interfaces to ensure it’s accessible for both bioinformaticians and clinicians.

## Introduction

Biological systems are inherently dynamic, and the temporal coordination of molecular processes is central to physiology, adaptation, and disease [1]. Recent advances in multi-omics technologies have enabled broad molecular profiling across metabolites, lipids, proteins, and cytokines [2]. However, most existing studies rely on sparsely sampled longitudinal designs, often with sampling intervals on the scale of days to months [3]. Such low temporal resolution limits the ability to infer dynamic regulatory relationships, particularly causal interactions that unfold on minute-to-hour time scales [4].

Time-series analysis provides a foundation for inferring causal relationships from temporal data [5]. Classical approaches such as Granger causality have been widely used to infer directionality by testing whether past values of one variable improve the prediction of another [6]. Extensions of these concepts have been developed for complex, multivariate, and nonlinear systems [7]. Methods such as convergent cross mapping and information-theoretic measures further attempt to capture causality in nonlinear systems [7]. More recently, causal discovery frameworks such as PCMCI have been introduced to identify causal relationships in high-dimensional time-series data while controlling for confounding and autocorrelation [8]. Despite these advances, existing methods often assume linear relationships, require strong stationarity assumptions, or become computationally intractable when applied to large-scale omics datasets [9].

In parallel, the emergence of high-throughput omics technologies has enabled unprecedented molecular coverage across multiple biological layers [2]. Large-scale human multi-omics studies have demonstrated the potential to uncover disease mechanisms and molecular networks [2, 10]. However, most multi-omics studies remain limited by low sampling frequency, restricting causal interpretation. To address this gap, dense longitudinal sampling strategies have begun to emerge. Microsampling technologies now allow minimally invasive, high-frequency sampling of human blood, enabling fine-grained temporal profiling of molecular dynamics [11]. Such a dataset provides a unique opportunity to infer dynamic and potentially causal relationships between molecular entities *in vivo*.

Here we present Lagged-Correlation Based Causal Inference (lagCI), a computational framework for inferring time-lagged causal relationships from high-frequency multi-omics time series. LagCI is based on our previous work with several improvements [11]. By systematically evaluating lagged correlations and assessing the robustness of lag profiles, lagCI distinguishes genuine delayed associations from spurious correlations. We validate lagCI using wearable physiological time-series data and demonstrate its ability to recover known causal relationships. We then apply lagCI to a dense human multi-omics dataset generated through microsampling, constructing a large-scale directed molecular causal network (http://lagomics.jaspershenlab.com/). The inferred network recapitulates known biological relationships and reveals novel molecular interactions, providing a systems-level view of temporal regulation in human physiology.

## Results

### Overview of lagCI

To uncover causal relationships between two biological time series, we developed lagCI, a computational algorithm that infers directionality based on the principle of lagged correlation [6]. Biological systems are inherently dynamic, and causal effects often manifest with a temporal delay [4]. That is, if variable X causally regulates variable Y, changes in X are typically followed by delayed responses in Y. This delay, known as lag time, causes the two time series to fluctuate asynchronously [12]. Capturing this temporal offset forms the foundation of the lagCI algorithm.

The lagCI algorithm begins by systematically shifting one time series relative to another across a user-defined lag window (Fig. 1a and Methods). For each shift, lagCI computes the Pearson correlation coefficient (ρ) and the corresponding *P-value*. This results in two vectors: *Lag_cor*, representing correlation coefficients across lags, and *Lag_P*, their associated significance values. To assess whether a strong correlation at a particular lag reflects a true temporal relationship or random fluctuation, lagCI fits the *Lag_cor* distribution to a Gaussian model, thereby generating predicted correlations (*Fitted_cor*) under the assumption of a null temporal structure (Fig. 1b). A quality score is then calculated as the absolute Spearman correlation between the observed *Lag_cor* and the *Fitted_cor* vectors. This metric quantifies the internal consistency of the lag-correlation profile, *i*.*e*., whether peak correlations are supported by a coherent trend across lag values rather than isolated outliers. Only time series pairs with high-quality scores are retained for downstream causal inference, ensuring that spurious or noisy correlations are filtered out (Methods). Finally, only the peak *Lag_cor* at non-zero lag (shift time) is considered as a causal relationship between two time series.

**Figure 1.**
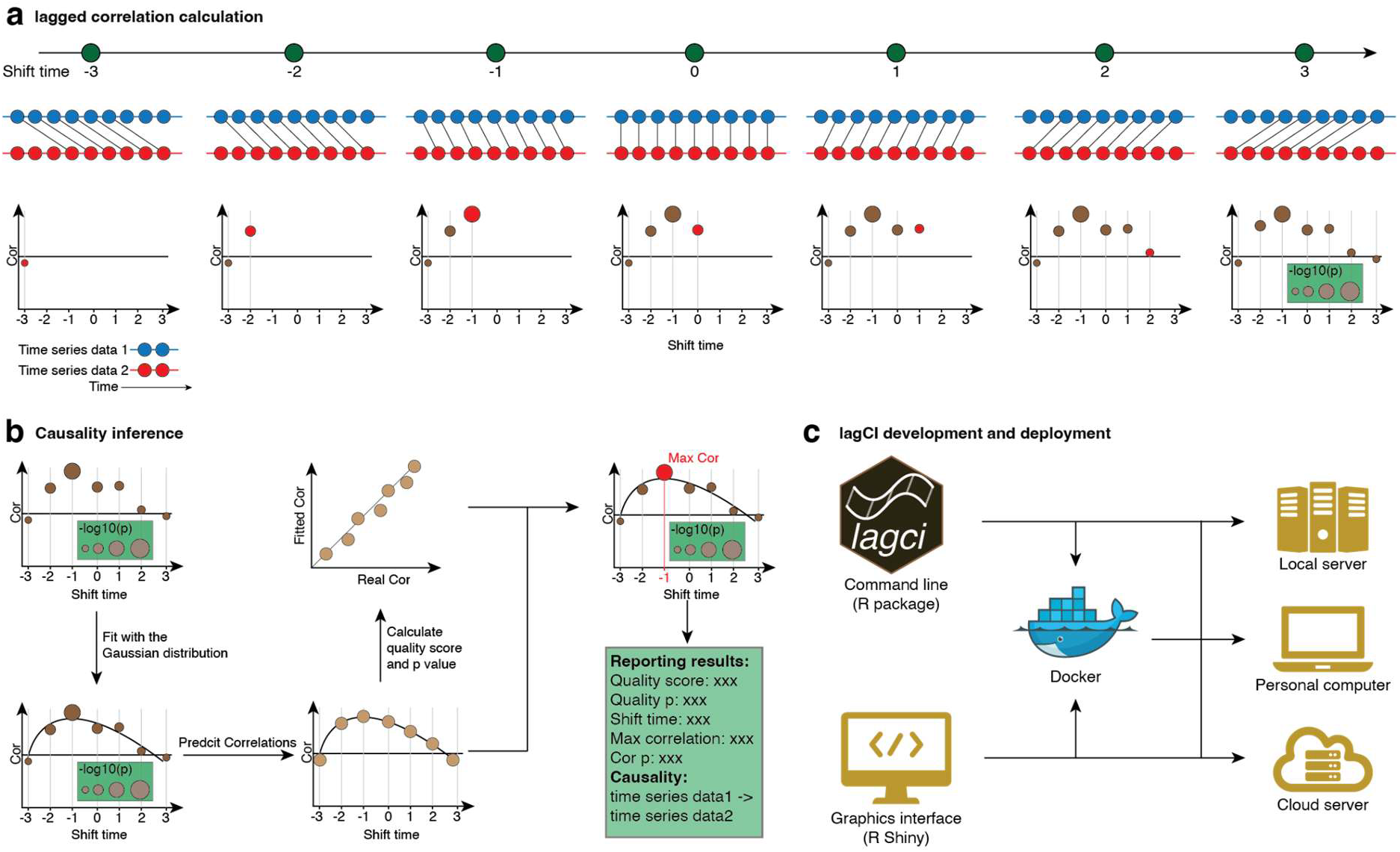
Overview of the lagCI algorithm for causal inference between time series. (a) Lagged correlation analysis. (b) Quality control and temporal causal relationship inference. (c) Software implementation and deployment.

To support widespread adoption, we implemented lagCI as a fully documented and open-source R package, designed for seamless integration into existing time-series or omics workflows. In addition to command-line functionality, lagCI offers a Shiny-based graphical user interface, enabling interactive visualization of lag profiles and inferred causal directionality. The entire pipeline is also containerized using Docker, facilitating reproducible deployment across local workstations, institutional clusters, and cloud platforms (Fig. 1c). For broader accessibility, we developed a publicly hosted version at https://lagcishiny.jaspershenlab.com, providing point-and-click functionality for researchers without programming expertise.

### Validation of lagCI using smartwatch data

To assess whether lagCI can reliably identify true causal relationships between biological time series, we applied the algorithm to a well-established physiological relationship: physical activity and heart rate. It is widely accepted that increases in physical activity (*e*.*g*., walking or running, quantified as steps) lead to elevated heart rates, a clear directional causal relationship [13]. This provided an ideal validation scenario.

We leveraged a publicly available wearable dataset from a previous study on early COVID-19 detection, which includes continuous step counts and heart rate measurements collected via smartwatches from 120 participants over approximately one month each (Methods and Supplementary Table 1) [14]. For each participant, we applied lagCI to evaluate the lagged correlation between step count and heart rate time series (Fig. 2a and Supplementary Table 2). Across all participants, we observed significant lagged correlations, indicating robust detection of the expected causal direction from steps to heart rate. Notably, the estimated lag times varied among individuals, clustering into three major groups based on the shift at which maximum correlation was observed. 53 participants showed peak correlation at a 1-minute lag, 64 participants at a 2-minute lag, and 2 participants at a 3-minute lag (Fig. 2b).

**Figure 2.**
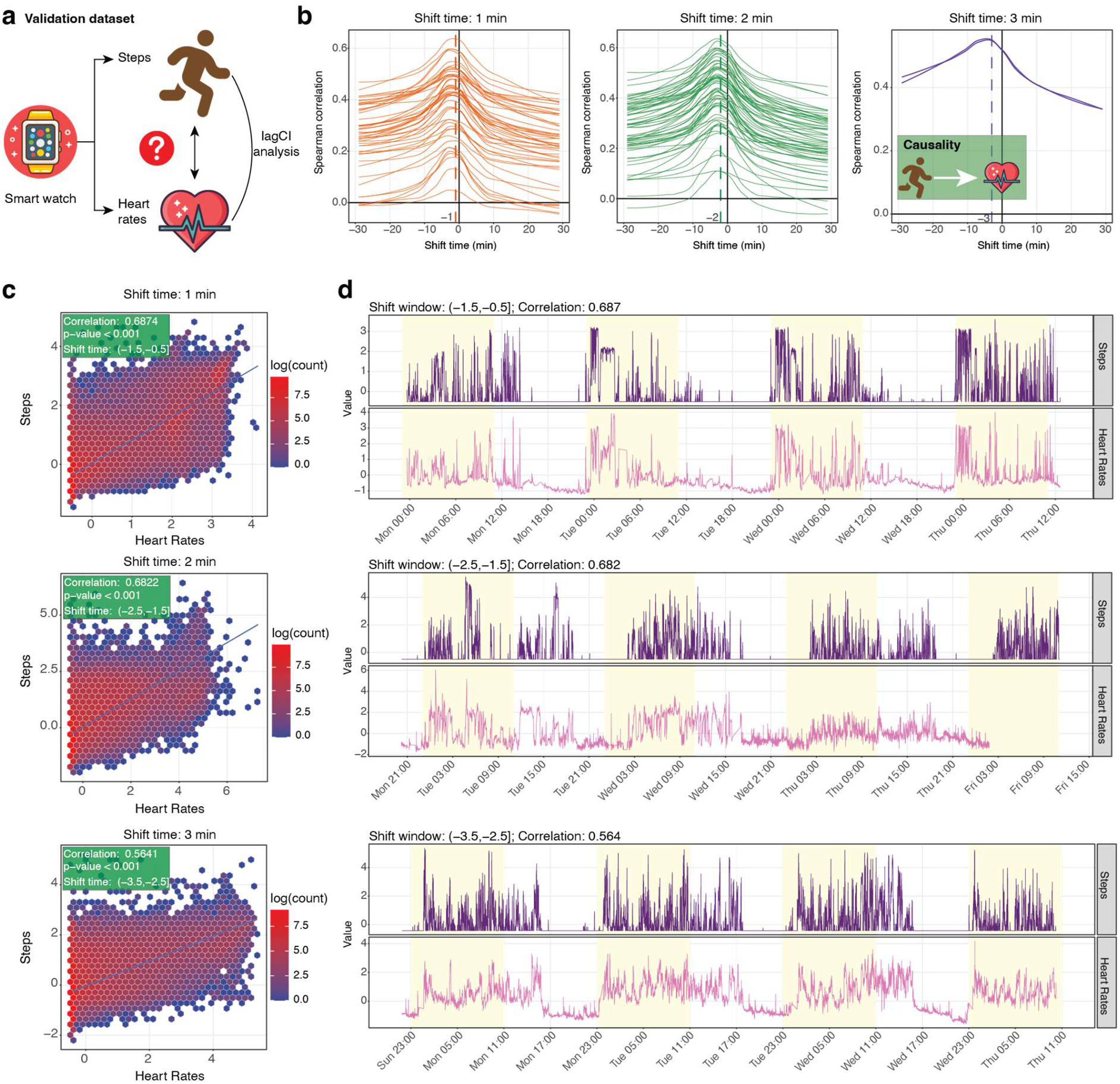
Validation of lagCI using step count and heart rate data from wearable smartwatches. (a) Schematic overview of the validation dataset. (b) Distribution of lagged correlation profiles across participants, stratified by peak correlation occurring at 1-minute (orange), 2-minute (green), or 3-minute (purple) lag. (c) Hexbin scatterplots of steps versus heart rate for one representative participant from each group. (d) Time series plots of step counts (purple) and heart rate (pink) for the same three representative participants.

To illustrate representative examples from each group, we selected one participant per cluster (Methods). In all cases, positive correlations between steps and shifted heart rates were evident (Fig. 2c), with Spearman correlation coefficients of 0.6874, 0.6822, and 0.5641 for 1-minute, 2-minute, and 3-minute shift times, respectively. These relationships are also visually apparent in time-aligned scatter and line plots, where bursts in step activity were consistently followed by heart rate increases (Fig. 2d).

These results demonstrate that lagCI accurately identifies the temporal directionality of this known physiological causal relationship at an individual level. The observed inter-individual differences in lag times are biologically plausible, likely reflecting variation in fitness level, cardiovascular responsiveness, or body mass index (BMI) among participants [15]. Demographic data were not available in the source dataset to further investigate these hypotheses [14]. Future studies with richer metadata will be necessary to explore the physiological drivers of such inter-individual variation in temporal coupling. In summary, this validation confirms that lagCI is a robust and generalizable tool for inferring temporal causal relationships from real-world, time series data, successfully capturing known dynamic biological relationships with individual-level resolution.

### LagCI uncovers extensive potential causal relationships among circulating molecules

To systematically uncover dynamic molecular regulation in humans, we next applied lagCI to infer potential causal relationships among circulating molecules. Biological processes *in vivo* operate across a wide range of time scales, from minutes to hours, necessitating high-resolution temporal sampling to resolve directionality and delays between molecular events [11, 16]. Conventional multi-omics studies are limited in this regard due to sparse sampling frequency. By contrast, microsampling-based multi-omics enables high-frequency longitudinal profiling of human blood [11]. We previously developed this platform to facilitate dense temporal sampling in humans [11]. Here, we leveraged a published dataset generated using this approach, in which a single participant collected fingertip blood samples every 2-3 hours over 7 consecutive days under free-living conditions, yielding 97 time points [11] (Fig. 3a-b). This dataset provides a unique opportunity to decode *in vivo* molecular causal relationships at high temporal resolution.

**Figure 3.**
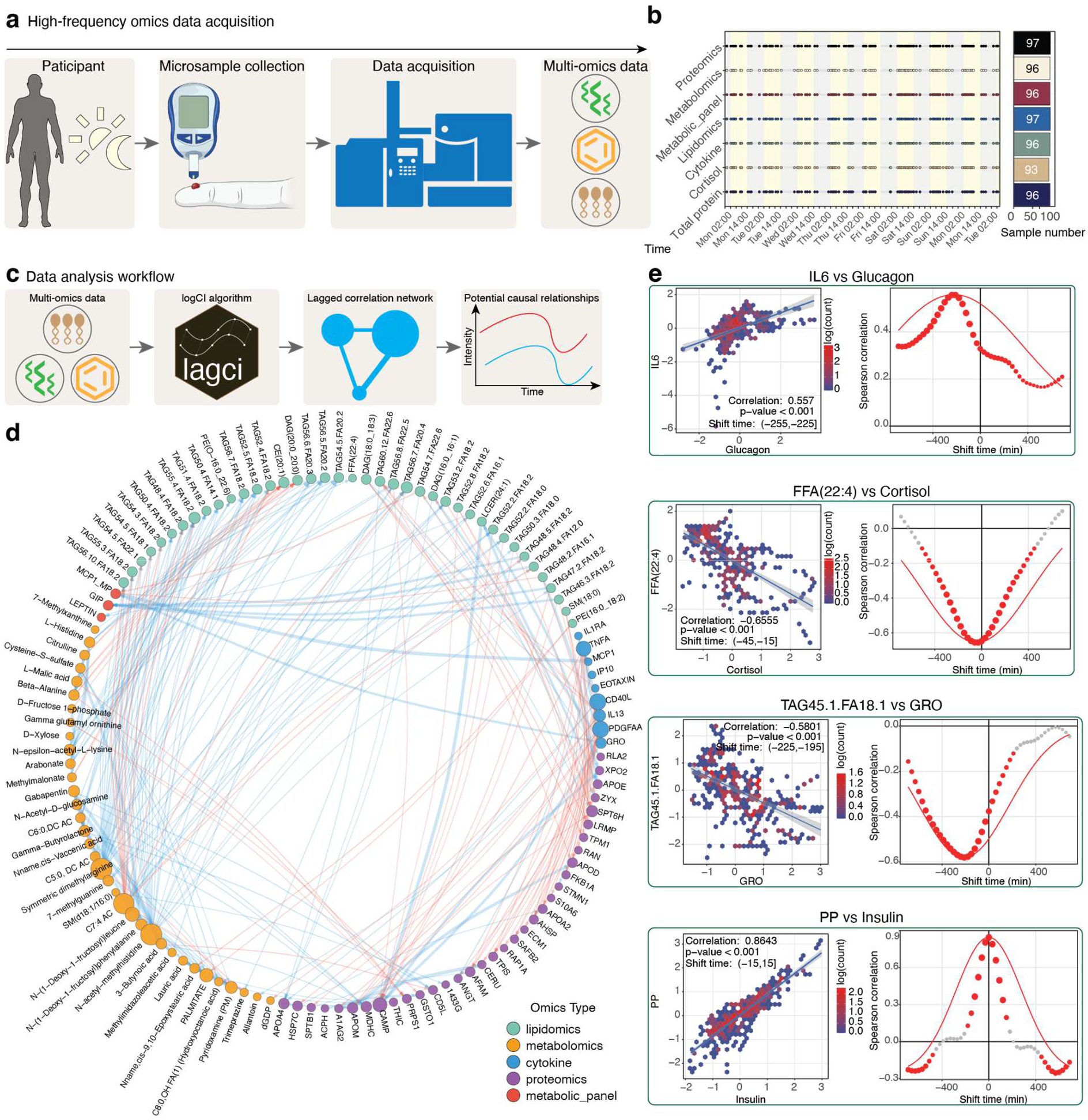
Inference of causal molecular relationships from high-frequency human multi-omics data. (a) Overview of the study design. (b) Distribution of multi-omics measurements across the sampling period. (c) Data analysis workflow. (d) Global molecular causal network inferred by lagCI (demo network). (e) Representative examples of biologically supported causal relationships identified by lagCI.

We analyzed this high-density multi-omics dataset using lagCI (Fig. 3c and Supplementary Table 2). Specifically, we performed exhaustive pairwise evaluation across 1,624 detected molecules, including 467 metabolites, 811 lipids, 292 proteins, 41 cytokines, and 13 hormones [11]. For each molecular pair, lagCI computed lagged correlation profiles, identified optimal delay times, and assigned quality scores reflecting signal strength and robustness. High-confidence causal links were defined using stringent criteria (absolute lagged correlation > 0.5, FDR-adjusted *p-value* < 0.05, and quality score > 0.8). After filtering, the final resulting causal network comprised 1,542 molecules connected by 157,489 directed edges, representing time-shifted relationships in which changes in one molecule precede changes in another (Fig. 3d).

This data-driven network revealed a rich and structured landscape of molecular interdependencies. The network exhibited a scale-free topology, with a small subset of highly connected hub molecules. The most connected node, DAG(18:2_22:5), showed a degree of 613, with lag times spanning -210 to 690 minutes, followed by DAG(14:0_18:2) (degree = 607) and DAG(16:0_18:2) (degree = 605), highlighting the central role of lipid species in coordinating metabolic and signaling processes [17]. Additional hub nodes included apolipoprotein E (degree = 338), gastric inhibitory polypeptide (GIP; degree = 320), and growth-regulated oncogene (GRO; degree = 82), suggesting active involvement of lipid, endocrine, and immune regulators in shaping temporal molecular dynamics [18]. These hubs likely represent key orchestrators of multi-omic transitions in circulation.

Importantly, lagCI recovered multiple well-established molecular relationships that are consistent with known physiological mechanisms (Fig. 3e). For example, an increase in IL-6 preceded a rise in glucagon by approximately 4 hours (lagged correlation = 0.557, BH-adjusted *p-value* = 0, shift time = -240 min), consistent with the role of IL-6 in modulating pancreatic α-cell activity and glucagon secretion [19]. Similarly, an increase in TAG(45:FA18:1) was followed by a decrease in the chemokine growth-regulated oncogene after approximately 3.5 hours (lagged correlation = -0.581, BH-adjusted *p-value* = 0, shift time = -210 min), in line with established links between lipid metabolism and inflammatory signaling [20]. We also observed that elevation of FFA(22:4) preceded a reduction in cortisol within approximately 30 minutes (lagged correlation = -0.655, BH-adjusted *p-value* = 0, shift time = -30 min), consistent with evidence that elevated free fatty acids can suppress hypothalamic-pituitary-adrenal axis activity [21]. Finally, insulin and pancreatic polypeptide displayed near-synchronous increases (lagged correlation = 0.8643, BH-adjusted *p-value* = 0, shift time = 0 min), reflecting their coordinated secretion in response to feeding and autonomic regulation [22].

Together, these results demonstrate that lagCI, when applied to dense microsampled human multi-omics data, can robustly infer directed molecular relationships that recapitulate known biology. The resulting causal network provides a systems-level framework for understanding how molecular signals propagate across multiple omic layers over time.

## Discussion

We present lagCI, a computational framework for inferring temporal directionality from biological time series by exploiting delayed correlation structure. Across two distinct settings, a smartwatch-based physiological validation task and a dense human blood multi-omics dataset, lagCI recovered time-ordered relationships that were consistent with known biology and identified a large number of previously unresolved molecular dependencies. Together with its open-source implementation in R, graphical Shiny interface, and containerized deployment, lagCI provides a practical and accessible framework for converting dense longitudinal data into biologically interpretable hypotheses about temporal ordering.

Many regulatory effects are not instantaneous [23], and biological responses often unfold over timescales ranging from minutes to hours. However, most commonly used methods assume synchronous relationships between variables [24]. lagCI was designed to address this gap. Rather than attempting to fully reconstruct mechanistic causation from observational data, lagCI asks a narrower and practically useful question: whether changes in one variable consistently precede changes in another with a reproducible lag structure. This framing is well-suited to dense longitudinal omics and wearable datasets, where temporal ordering can provide strong clues about regulatory organization even when intervention data are unavailable.

The smartwatch analysis illustrates this point clearly. lagCI recovered the expected direction from physical activity to heart rate across individuals and resolved subject-specific lag times on the order of minutes [25]. This result serves as an important validation because the underlying physiological relationship is well established, while the observed inter-individual variation is itself biologically plausible. Differences in cardiovascular fitness, autonomic responsiveness, metabolic state, or other personal factors could all contribute to distinct temporal delays between motion and heart-rate response [26]. More broadly, this result highlights one strength of lagCI: it operates at the individual level, enabling detection of individual-specific temporal structure rather than collapsing all dynamics into cohort averages.

The application to dense human multi-omics data further demonstrates the potential of lagCI in systems biology. Using high-frequency microsampled blood measurements [11], lagCI generated a directed molecular network spanning metabolites, lipids, proteins, cytokines, and hormones. This network recapitulated multiple known relationships, including interactions linking inflammatory cytokines, metabolic hormones, and endocrine regulators, lending confidence to the biological validity of the inferred causal relationships.

An important feature of lagCI is that it does not rely solely on the maximum lagged correlation. Instead, it evaluates the overall shape of the lag-correlation profile and prioritizes relationships with coherent, peak-supported structure. This quality-control step is intended to distinguish meaningful delayed relationships from isolated peaks caused by random alignment, noise, or local fluctuations. In practice, this proved valuable in both validation and application settings, where a large number of possible pairwise comparisons could otherwise generate unstable or difficult-to-interpret results. By combining lag scanning and profile-level quality assessment, lagCI provides a transparent and scalable analysis strategy that remains straightforward to interpret. The availability of the same workflow through command-line execution, graphical exploration, and Dockerized deployment further lowers the barrier to use across a wide range of computational environments.

At the same time, several limitations should be emphasized. First, lagCI infers temporal directionality, not mechanistic causation in the strict experimental sense. A delayed X→Y relationship may reflect direct regulation, indirect mediation through unmeasured intermediates, shared upstream drivers, feedback loops, or structured environmental inputs. As with other observational causal discovery approaches, temporal precedence strengthens biological plausibility but does not by itself establish a mechanism. Second, the method depends on data quality and preprocessing. Irregular sampling, missingness, smoothing, interpolation, and detrending choices can all affect the estimated lag structure. Although our framework restricts inference to aligned observations and incorporates profile-level filtering, preprocessing remains an important determinant of reliability. Third, pairwise lagged correlation cannot fully account for confounding in complex biological systems. In dense molecular networks, many variables are strongly interdependent, and some inferred edges may represent indirect rather than direct relationships. Fourth, the multi-omics application presented here was derived from a single individual. Although this design is valuable for demonstrating feasibility and revealing high-resolution biology, it does not establish the population-level reproducibility of all inferred relationships.

These limitations point naturally to future directions. Extending lagCI beyond pairwise analysis toward multivariate or conditional frameworks would help distinguish direct from indirect temporal dependencies and better address confounding. Incorporating nonlinear similarity measures or information-theoretic dependence may also improve sensitivity to relationships that are not well captured by linear correlation. At the study-design level, applying lagCI to larger cohorts with repeated dense sampling will be essential for identifying which lagged relationships are personal, which are conserved, and how temporal coupling varies with physiology, disease, or intervention. Finally, combining lagCI with perturbational or interventional studies could provide a powerful route from time-ordered association to experimentally supported mechanism.

In summary, lagCI provides a statistically grounded and user-accessible framework for inferring temporal directionality from biological time series. By coupling lag-profile evaluation with scalable software and dense longitudinal data, it enables the recovery of known biology and the discovery of previously unresolved time-ordered molecular relationships. We anticipate that lagCI will be broadly useful for analyzing wearable, physiological, and multi-omics time series, and that it will serve as a foundation for future efforts in causal discovery from high-resolution longitudinal biology.

## Methods

### Overview of lagCI

We implemented a computational framework for calculating lagged correlations between time-series data using the *lagci* package. This method is designed to analyze the temporal relationships between omics and wearable device data. The lagCi is based on our previous work [11] with several improvements.

Prior to correlation analysis, all input time-series data underwent standardization by z-score normalization to ensure comparable scales between different data types. Time-series alignment was performed to establish a common temporal reference frame by determining the overlapping time range between datasets. The aligned time series were interpolated onto a regular time grid using either linear or constant interpolation methods. Missing data points were handled using rule-based extrapolation at time series boundaries to minimize data loss.

The core lagged correlation analysis utilized the cross-correlation function (CCF) to compute correlations across different time lags [27]. Both Spearman and Pearson correlation methods were supported, with rank transformation applied for Spearman correlations. Statistical significance of correlations was evaluated using normal approximation.

Optional data smoothing was implemented using locally weighted scatterplot smoothing (LOESS) regression. The smoothing parameters included span values and polynomial degree, which could be optimized based on data characteristics.

The quality of lagged correlation results was evaluated using a specialized scoring system that fits LOESS curves to correlation data and performs peak detection. The method normalized correlations to handle both positive and negative relationships and computed quality scores based on Spearman correlation between fitted peaks and observed data.

Results were returned as structured objects containing correlation coefficients, *p-values*, shift times, and indices for maximum and global (zero-lag) correlations. Visualization functions provided scatter plots, alignment plots, and time-series plots to facilitate the interpretation of temporal relationships in the data.

The more detailed methods about lagCI have been provided in the Supplementary Note.

### Time alignment & missingness

To reduce edge artifacts, no extrapolation beyond observed boundaries was applied in the final analyses; instead, lag scans were restricted to the overlapping window across series. Internal gaps shorter than a preset threshold were optionally imputed (linear or last-observation-carried-forward), whereas longer gaps remained missing and were excluded pairwise during correlation evaluation. This policy follows the practical guidance in the tutorial’s data-processing chapter.

### Smoothing policy

LOESS smoothing was used only as an optional visualization aid. All statistical inference, including significance testing, FDR control, and peak-lag determination, was performed on the raw, unsmoothed series to preserve the serial-dependence structure essential for identifying temporal lags.

### Robustness analyses

We systematically varied Δt, τ_max, the correlation metric (Pearson *vs*. Spearman), and interpolation strategy. Stability was summarized by the agreement of τ^ sign/magnitude across settings and by **peak sharpness** (top-1 vs. top-2 ∣r∣gap).

### Deployment of lagCI and *lagcishiny*

We have deployed lagCI and *lagcishiny* in multiple formats. Both tools are implemented as R packages and can be installed directly from GitHub: the command-line version (lagCI) is available at https://github.com/jaspershen-lab/lagci, and the graphical interface (*lagcishiny*) at https://github.com/jaspershen-lab/lagcishiny.

To ensure seamless cross-platform compatibility and reproducibility, we also provide Dockerized versions of both tools, available via Docker Hub at https://hub.docker.com/r/jaspershenlab/lagci. For users who prefer not to install any software, a fully hosted and interactive online instance of *lagcishiny* is maintained at https://lagcishiny.jaspershenlab.com, offering immediate access through a web browser. Comprehensive installation guides, usage tutorials, and deployment instructions for all versions are available at https://www.shen-lab.org/lagci-tutorial.

### Demo datasets

#### Wearable dataset

The wearable dataset used to validate the lagCI algorithm was obtained from a previously published study on early detection of COVID-19 using smartwatch data [14]. Briefly, the dataset includes continuous time-series recordings of physiological signals, primarily step count and heart rate. The publicly available dataset contains wearable data from 120 individuals in total, including 32 COVID-19 positive cases, 15 with non-COVID respiratory illnesses, and 73 contemporaneous healthy controls. The data were captured using consumer-grade wearable devices such as Fitbit and Apple Watch, spanning periods both before and after symptom onset.

To validate the ability of lagCI to detect temporally shifted relationships, we selected representative pairs of physiological variables, namely, heart rate and step count, and applied lagCI to infer whether one signal anticipates changes in the other. Only individuals with a time series overlap between heart rate and step count were included in this analysis.

#### Dense human multi-omics dataset

The dense human multi-omics dataset was obtained from a previously published study on high-frequency microsampling for lifestyle-associated health profiling [11]. Briefly, a single participant was enrolled. To enable dense longitudinal sampling while maintaining participant compliance, a self-collection protocol was implemented: the participant performed finger-prick microsampling approximately once per hour during waking hours and once every two hours during overnight periods, over a span of seven consecutive days. Immediately after collection, each sample was placed on dry ice and shipped daily to the laboratory for processing. A total of 97 microsamples were collected and subjected to in-depth multi-omics profiling, including: (1) untargeted proteomics, (2) untargeted metabolomics, (3) semi-targeted lipidomics, and (4) targeted assays for cytokines, hormones, total protein, and cortisol.

### General statistics

Multiple hypothesis testing was corrected using the False Discovery Rate (FDR) method. Icons used in figure illustrations were sourced from iconfont.cn and are licensed under the MIT license for non-commercial use (https://pub.dev/packages/iconfont/license).

## Data availability

All datasets analyzed in this study were derived from previously published studies [11, 14]. No new datasets were generated as part of this work.

## Code availability

All software development, data processing, and statistical analyses were performed using R (version 4.4.1) and associated packages (Supplementary Note). The code for the lagCI algorithm and its graphical interface, *lagcishiny*, is available at https://github.com/jaspershen-lab/lagci and https://github.com/jaspershen-lab/lagcishiny, respectively.

## Author contributions

XS conceived the project and provided overall supervision. XS, SB, YG, and ZQ jointly designed the methodology and developed the software. The Docker-based implementation was contributed by YL and SB. Documentation and user tutorials were prepared by SB, YG, ZQ, and XS. Case study analyses were performed by YG, SB, YW, ZQ, and XS, who also generated all figures. The manuscript was written by XS, YG, and SB, with input from all authors. All authors reviewed and approved the final version of the manuscript.

## Additional information

Correspondence and requests for materials should be addressed to XS.

## Key points

1. We developed Lagged-Correlation Based Causal Inference (lagCI), a computational framework designed to identify time-lagged associations by combining comprehensive lag-correlation profiling with a robust statistical filtering scheme, which can identify the potential causal relationships between molecules in multi-omics data.
2. We applied lagCI to dense human multi-omics data and successfully recovered well-known physiological relationships while constructing a large-scale directed molecular causal network of over 157,000 predicted interactions, demonstrating its capacity to uncover dynamic molecular regulation and resolve individual-specific lag times.
3. The framework is implemented as an open-source R package with a Shiny-based graphical user interface and Dockerized deployment options, providing researchers with an accessible, transparent, and scalable analysis strategy for converting dense longitudinal data into biologically interpretable hypotheses.

## Biographical Note

Xiaotao Shen is an Assistant Professor at Lee Kong Chian School of Medicine, Nanyang Technological University, Singapore. His research focuses on developing bioinformatics algorithms for multi-omics data integration and applying them to precision medicine.

