## Supplementary Information for "LagCI Enables Inference of Temporal Causal Relationships from Dense Multi-Omic Time Series"

**Supplementary Note**
**lagCI software**
*Software implementation.* We developed lagci, an R package for time-resolved association analysis between two longitudinal signals, particularly for wearable and omics data. The package was implemented in R (version $\geq$ 4.1) and distributed as an MIT-licensed package. Core imports include stats, lubridate, BiocParallel, ggplot2, dplyr, massdataset, and related visualization and utility packages, enabling time handling, interpolation, correlation analysis, parallel computation, and result visualization within a unified workflow

The package exposes a compact set of primary analysis functions, including calculate_lagged_correlation(), smooth_data(), fit_smoothing_parameters(), and evaluate_lagged_cor(), together with accessor and plotting functions for downstream interpretation of lagged correlation profiles and aligned time-series relationships

*Data model and result object.* Lagged correlation results are stored in an S4 class, lagged_cor_result, which retains the original input vectors (x, y), their timestamps (time1, time2), the full lag-resolved correlation profile (all_cor), corresponding P values (all_cor_p), lag interval labels (shift_time), the index of the maximum absolute correlation (which_max_idx), the index corresponding to the zero-lag correlation (which_global_idx), the maximum correlation value (max_cor), the zero-lag correlation value (global_cor), and a structured parameter record (tidymass_parameter) describing the analysis settings used to generate the result. This object design allows the full lagged-correlation landscape, rather than only a single summary statistic, to be preserved for downstream scoring and visualization.

*Optional preprocessing and smoothing of time-series data.* To reduce high-frequency noise prior to downstream analysis, lagci implements an optional LOESS-based smoothing workflow. In smooth_data(), the observed signal is modeled as a function of numeric time using loess(x ~ as.numeric(time)), with user-specified span and polynomial degree, and the fitted values are returned together with the original data. An optional plotting mode overlays the original and smoothed trajectories for visual inspection of smoothing behavior.

To assist parameter selection, the package also provides fit_smoothing_parameters(), which performs a grid search over candidate LOESS span and degree values. The input series is first sorted by time, and the first one-third of the observations is used as a fitting subset. After excluding missing values and values $\leq 0$, LOESS models are fit across the candidate parameter grid in parallel using BiocParallel. Each model is then used to predict the full original series, and the squared Pearson correlation ($R^2$) between observed and predicted values is calculated. The final smoothing parameters are chosen as the combination whose $R^2$ is closest to 0.6, rather than the absolute maximum $R^2$, thereby favoring moderate smoothing over potentially overfit trajectories. By default, span values from 0.1 to 0.5 in increments of 0.01 and polynomial degrees 1 or 2 are evaluated.

*Calculation of lagged correlation.* The main analysis function, calculate_lagged_correlation(), estimates the correlation between two time-series signals across a user-defined lag window. Inputs include two numeric vectors (x and y), their timestamp vectors (time1 and time2), the total lag tolerance (time_tol, in hours), the lag step size (step, in hours), the minimum number of matched observations required for a valid analysis, the alignment method, and the correlation method (spearman or pearson).

Before alignment, both input vectors are converted to numeric and standardized using z-score scaling. The analysis is restricted to the overlapping time interval defined by the later of the two start times and the earlier of the two end times. A regular time grid is then generated across this shared interval at the requested temporal resolution (step × 60 minutes). Each input series is interpolated onto this common grid using stats::approx(), with either linear or constant interpolation and with edge extrapolation enabled (rule = 2) to avoid missing values at the boundaries of the overlap interval

For Spearman analysis, the aligned values are rank-transformed prior to correlation calculation; for Pearson analysis, the aligned values are used directly. Complete paired observations are retained, and analyses are discarded if the number of effective matched samples is smaller than either 2 or the user-specified min_matched_sample threshold. The maximum lag index is computed as round(time_tol / step) - 1 and capped at n_eff - 1, where n_eff is the number of complete aligned observations. Lagged correlations are then calculated using the discrete cross-correlation function (stats::ccf) without plotting.

From the ccf output, the package extracts the full lag-dependent correlation profile (all_cor) and computes two-sided P values by normal approximation under a null centered at zero, using a standard deviation of 1 / sqrt(n_used), where n_used is the number of observations used in the cross-correlation calculation. The lag axis is represented as consecutive shift intervals centered on the integer lag positions and expressed in minutes. The package records both the lag with the maximum absolute correlation and the zero-lag position, allowing users to distinguish delayed associations from synchronous associations within the same result object

*Evaluation of lagged-correlation profile shape.* Beyond identifying the largest lagged association, lagci implements an additional profile-level scoring procedure in evaluate_lagged_cor(). This function takes a lagged_cor_result object and evaluates whether the lag-correlation curve exhibits a coherent peak-like structure rather than a noisy or fragmented profile. To do this, the function first converts the lag-interval labels into numeric shift times by taking the midpoint of each lag interval. Missing correlation values are set to zero, and a LOESS curve is fit to the lag-correlation profile as a function of shift time. The smoothing span is set adaptively to 3 / length(cor) with a lower bound of 0.2, and predictions are generated on a 1-minute grid spanning the lag range.

The smoothed lag-correlation curve is then decomposed into positive and negative candidate envelopes. For the positive component, the curve is shifted upward if needed so that all values are non-negative, the dominant positive peak is identified, and a Gaussian peak shape is reconstructed using fitted location, width, and area parameters. The similarity between the reconstructed peak and the positive envelope is quantified using Spearman correlation. The same procedure is repeated for the negative component after sign inversion. The final score is defined as the larger of the positive and negative peak-fit scores, provided that the original lag-correlation profile actually contains a positive or negative extremum of the corresponding sign. When plotting is enabled, the function returns a visualization showing the smoothed lag-correlation profile, the fitted peak, the significance of observed lag bins, and the zero-shift reference line.

*Visualization and interpretation of lagged associations.* The package includes dedicated plotting utilities to visualize lagged relationships from complementary perspectives. lagged_alignment_plot() displays the temporal alignment between the two input series at either the zero-lag ("global") position or the lag corresponding to the maximum absolute correlation, with options to display matched points, integrated averages, and connection lines between aligned observations. lagged_scatter_plot() summarizes the relationship between the two signals at a selected lag by plotting matched values as either a scatter plot or a hexbin plot and overlays a fitted linear trend for visual interpretation of the lag-specific association strength.

*Parallelization and reproducibility.* Where applicable, computationally intensive steps are parallelized using BiocParallel. In fit_smoothing_parameters(), parallel backends are chosen in an operating-system-aware manner: SnowParam is used on Windows and MulticoreParam on non-Windows systems. A random seed can be supplied to ensure reproducible parameter selection across runs. In calculate_lagged_correlation(), analysis parameters are recorded in a tidymass_parameter object and stored in the returned lagged_cor_result, preserving the exact settings used for lag tolerance, step size, sample threshold, thread count, and correlation method.

**R packages used in the study**

R version 4.5.2 (2025-10-31)
Platform: aarch64-apple-darwin20
Running under: macOS Tahoe 26.3.1

Matrix products: default
BLAS:
/System/Library/Frameworks/Accelerate.framework/Versions/A/Frameworks/vecLib.framework/Versions/A/libBLAS.dylib
LAPACK: /Library/Frameworks/R.framework/Versions/4.5-arm64/Resources/lib/libRlapack.dylib; LAPACK version 3.12.1

locale:
[1] en_US.UTF-8/en_US.UTF-8/en_US.UTF-8/C/en_US.UTF-8/en_US.UTF-8

time zone: Asia/Singapore
tzcode source: internal

attached base packages:
[1] tcltk stats graphics grDevices utils datasets methods base

other attached packages:
[1] vegan_2.7-1 permute_0.9-8 scales_1.4.0 pheatmap_1.0.13 Mfuzz_2.68.0 DynDoc_1.86.0
[7] widgetTools_1.86.0 e1071_1.7-16 Biobase_2.68.0 BiocGenerics_0.54.0 generics_0.1.4 lubridate_1.9.4
[13] forcats_1.0.0 stringr_1.5.1 dplyr_1.1.4 purrr_1.1.0 readr_2.1.5 tidyr_1.3.1

```
[19] tibble_3.3.0       ggplot2_4.0.2       tidyverse_2.0.0    magrittr_2.0.3     r4projects_0.1.7

loaded via a namespace (and not attached):
  [1] DBI_1.2.3                tmaptools_3.3            remotes_2.5.0            rlang_1.1.6
  [5] ade4_1.7-23              clue_0.3-66              GetoptLong_1.0.5         matrixStats_1.5.0
  [9] compiler_4.5.2           mgcv_1.9-3               reshape2_1.4.4           png_0.1-8
 [13] systemfonts_1.2.3        vctrs_0.6.5              rvest_1.0.4              pkgconfig_2.0.3
 [17] shape_1.4.6.1            crayon_1.5.3             XVector_0.48.0           lwgeom_0.2-14
 [21] labeling_0.4.3           utf8_1.2.6               tzdb_0.5.0               UCSC.utils_1.4.0
 [25] ragg_1.4.0               bit_4.6.0                GenomeInfoDb_1.44.2      jsonlite_2.0.0
 [29] biomformat_1.36.0        rhdf5filters_1.20.0      DelayedArray_0.34.1      Rhdf5lib_1.30.0
 [33] parallel_4.5.2           cluster_2.1.8.1          R6_2.6.1                 stringi_1.8.7
 [37] RColorBrewer_1.1-3       GenomicRanges_1.60.0     stars_0.6-8              Rcpp_1.1.0
 [41] SummarizedExperiment_1.38.1 iterators_1.0.14      IRanges_2.42.0           igraph_2.1.4
 [45] splines_4.5.2            rentrez_1.2.4            Matrix_1.7-4             timechange_0.3.0
 [49] tidyselect_1.2.1         rstudioapi_0.17.1        stringdist_0.9.15        dichromat_2.0-0.1
 [53] abind_1.4-8              doParallel_1.0.17        codetools_0.2-20         plyr_1.8.9
 [57] lattice_0.22-7           withr_3.0.2              S7_0.2.0                 survival_3.8-3
 [61] sf_1.0-21                units_0.8-7              proxy_0.4-27             zip_2.3.3
 [65] xml2_1.4.0               Biostrings_2.76.0        circlize_0.4.16          phyloseq_1.52.0
 [69] pillar_1.11.0            MatrixGenerics_1.20.0    tkWidgets_1.86.0         KernSmooth_2.23-
26
 [73] foreach_1.5.2            stats4_4.5.2             vroom_1.6.5              S4Vectors_0.48.0
 [77] hms_1.1.3                class_7.3-23             glue_1.8.0               tools_4.5.2
 [81] data.table_1.17.8        ggsignif_0.6.4           openxlsx_4.2.8           XML_3.99-0.19
 [85] rhdf5_2.52.1             grid_4.5.2               ape_5.8-1                massdataset_0.99.3
 [89] colorspace_2.1-1         nlme_3.1-168             GenomeInfoDbData_1.2.14  cli_3.6.5
 [93] textshaping_1.0.1        S4Arrays_1.8.1           ComplexHeatmap_2.24.1    gtable_0.3.6
 [97] digest_0.6.37            classInt_0.4-11          SparseArray_1.8.1        rjson_0.2.23
[101] farver_2.1.2             multtest_2.64.0          lifecycle_1.0.4          httr_1.4.7
[105] GlobalOptions_0.1.2      MASS_7.3-65              bit64_4.6.0-1
```

**Supplementary Tables**

**Supplementary Table 1. Wearable individuals informations**

| id | lag_center_min | max_cor | global_cor | quality_score |
|---|---|---|---|---|
| A06L7KF | -1 | 0.514741987 | 0.502253149 | 0.979544126 |
| A0822M0 | -2 | 0.582396984 | 0.485506655 | 0.972472238 |
| A0KX894 | -2 | 0.398097241 | 0.347151115 | 0.947340736 |

| | | | | |
|---|---|---|---|---|
| A0L9BM2 | -1 | 0.485373215 | 0.447154144 | 0.984862653 |
| A0N9NV4 | -1 | 0.472216692 | 0.449269941 | 0.949444769 |
| A0NVTRV | -2 | 0.121059266 | 0.105145817 | 0.9471654 |
| A0VFT1N | -1 | 0.460630254 | 0.434315022 | 0.985797779 |
| A11SQQN | -1 | 0.344435002 | 0.308962928 | 0.966510812 |
| A11V1FH | -2 | 0.498078735 | 0.455573362 | 0.972998247 |
| A17YCA2 | -2 | 0.577368195 | 0.52811414 | 0.981297487 |
| A1K5DRI | -1 | 0.553551696 | 0.536278653 | 0.978667446 |
| A1ZJ41O | -2 | 0.448312398 | 0.387009771 | 0.966101695 |
| A2D7K4A | -2 | 0.413277332 | 0.361124323 | 0.981180596 |
| A2P3LTM | -2 | 0.503615637 | 0.469837532 | 0.976271186 |
| A2XFW2N | -1 | 0.067810854 | 0.054861135 | 0.87182934 |
| A35BJNV | -2 | 0.422317702 | 0.372048175 | 0.987960257 |
| A36HR6Y | -1 | 0.245567034 | 0.216737504 | 0.959964933 |
| A3OU183 | -1 | 0.272910194 | 0.263743062 | 0.981940386 |
| A45F9E6 | -2 | 0.343137024 | 0.273749916 | 0.974284044 |
| A4E0D03 | -2 | 0.509205817 | 0.436776998 | 0.966569258 |

| | | | | |
|---|---|---|---|---|
| A4G0044 | -1 | 0.485523242 | 0.464415049 | 0.957276447 |
| A4H7SNF | -2 | 0.534571545 | 0.471155674 | 0.989830508 |
| A5XL2IC | -1 | 0.584941477 | 0.471022652 | 0.980713033 |
| A65HVGP | -1 | 0.595351402 | 0.554042531 | 0.998129749 |
| A6BUI4N | -2 | 0.637700792 | 0.577742306 | 0.990414962 |
| A6GEBIK | -2 | 0.428687275 | 0.348037262 | 0.977206312 |
| A7EAWA7 | -2 | 0.446354098 | 0.39766012 | 0.97936879 |
| A7EM0B6 | -2 | 0.502214271 | 0.411399496 | 0.983167738 |
| A8CBEJZ | -2 | 0.625281459 | 0.567389093 | 0.986732905 |
| A8QLAB0 | -1 | 0.437231937 | 0.417501115 | 0.959848042 |
| A91HEZV | -1 | 0.63978545 | 0.599289847 | 0.9735827 |
| A99ZKKW | -2 | 0.321356973 | 0.304567968 | 0.978550555 |
| A9ZG5GR | -2 | 0.521617278 | 0.480833112 | 0.952308591 |
| AA0HAI1_1 | -1 | 0.253945985 | 0.206796394 | 0.975803624 |
| AA0HAI1_2 | -1 | 0.271152393 | 0.187982306 | 0.960081823 |
| AA0HAI1_3 | -2 | 0.252170332 | 0.15481091 | 0.952542373 |
| AA2KP1S | -2 | 0.473157719 | 0.394364625 | 0.977673875 |

| | | | | |
|---|---|---|---|---|
| AAF9ACE | -2 | 0.557013464 | 0.503441284 | 0.973056692 |
| AAGTWZK | -1 | 0.546770191 | 0.519309227 | 0.960900058 |
| AAXAA7Z | -2 | 0.595243042 | 0.524391983 | 0.986732905 |
| AD77K91 | -1 | 0.340243735 | 0.330220448 | 0.961075395 |
| AE0MQ94 | -1 | 0.589414946 | 0.576956914 | 0.992986558 |
| AE2B3RH | -1 | 0.359406936 | 0.327185931 | 0.969725307 |
| AEOBCFJ | -2 | 0.612754739 | 0.569373257 | 0.988077148 |
| AEOHH30 | -2 | 0.471835666 | 0.401614912 | 0.971186441 |
| AEZDKVO | -2 | 0.364366859 | 0.287348711 | 0.988603156 |
| AF3J1YC | -3 | 0.564116438 | 0.511816478 | 0.980420807 |
| AF3MXM1 | -2 | 0.559361233 | 0.503744474 | 0.963588545 |
| AF8R0I6 | -1 | 0.542696122 | 0.504583631 | 0.979485681 |
| AFEFA29 | -1 | 0.408998262 | 0.375315335 | 0.975920514 |
| AFHOHOM | -2 | 0.281518701 | 0.2450562 | 0.974810053 |
| AFPB8J2 | -1 | 0.522273149 | 0.456600149 | 0.950730567 |
| AFVAEC7 | -1 | 0.463728939 | 0.446414631 | 0.980362361 |
| AFYLHG4 | -1 | 0.312304636 | 0.293674424 | 0.967036821 |

| | | | | |
|---|---|---|---|---|
| AGA8XUN | -1 | 0.458086797 | 0.444793987 | 0.990648743 |
| AGKI03N | -2 | 0.315039461 | 0.269500814 | 0.994389246 |
| AGYQJEL | -2 | 0.660037039 | 0.616228807 | 0.987258913 |
| AHP25OJ | -1 | 0.286551221 | 0.255221751 | 0.955289305 |
| AHYIJDV | -2 | 0.419180317 | 0.375273526 | 0.982057276 |
| AHYR55C | -1 | 0.177506006 | 0.176803682 | 0.921040327 |
| AIFDJZB | -2 | 0.604119303 | 0.538629635 | 0.98351841 |
| AJ0DKQ3 | -2 | 0.622993018 | 0.575533924 | 0.978082992 |
| AJ7TSV9 | -2 | 0.325017833 | 0.275101307 | 0.976154296 |
| AJMQUVV | -2 | 0.499427175 | 0.46052824 | 0.974926943 |
| AJWW3IY | -1 | 0.456155678 | 0.41763402 | 0.973641146 |
| AK7YRBU | -2 | 0.334108878 | 0.275564989 | 0.99333723 |
| AKTGD8X | -1 | 0.527568937 | 0.515584125 | 0.989596727 |
| AKV66US | -2 | 0.277142019 | 0.261009731 | 0.960724722 |
| AKXN5ZZ | -2 | 0.49183285 | 0.409731073 | 0.964640561 |
| AL3KT5B | -2 | 0.629191157 | 0.581068455 | 0.954412624 |
| AL48GP3 | -2 | 0.620901425 | 0.558663961 | 0.979485681 |

| | | | | |
|---|---|---|---|---|
| ALKAXMZ | -1 | 0.517171388 | 0.495257143 | 0.977556984 |
| ALZDAVZ | -2 | 0.462608727 | 0.401658088 | 0.990414962 |
| AMQUHOQ | -1 | 0.448833835 | 0.434106368 | 0.992109877 |
| AMV7EQF | -2 | 0.414301078 | 0.39226991 | 0.994038574 |
| AO20DS4 | -2 | 0.423526964 | 0.366200867 | 0.958679135 |
| AOB9SON | -2 | 0.53561187 | 0.510901856 | 0.984862653 |
| AOGFRXL | -2 | 0.682223452 | 0.601570469 | 0.981239041 |
| AOQA85X | -1 | 0.518782635 | 0.503499699 | 0.985388662 |
| AOS4BSJ | -1 | 0.560494609 | 0.541989125 | 0.990005845 |
| AOYM4KG | -1 | 0.523342934 | 0.475957541 | 0.974868498 |
| APDJ1QP | -3 | 0.558216064 | 0.518348881 | 0.973641146 |
| APGIB2T | -1 | 0.469264785 | 0.454805838 | 0.984102864 |
| APHNRSV | -1 | 0.260520512 | 0.199534591 | 0.983635301 |
| AQ25Y0L | -2 | 0.463993755 | 0.394957418 | 0.974400935 |
| AQ4TMLV | -2 | 0.519334185 | 0.486000295 | 0.975160725 |
| AQC0L71 | -1 | 0.533341467 | 0.500885766 | 0.985856224 |
| AQR8ZSS | -1 | 0.530889193 | 0.525884183 | 0.989713618 |

| | | | | |
|---|---|---|---|---|
| AR4FPCC | -1 | 0.627674859 | 0.595074311 | 0.987784921 |
| ARFYLMK | -2 | 0.504756888 | 0.462558628 | 0.988836937 |
| ARR2IKE | -2 | 0.366838144 | 0.325690486 | 0.952600818 |
| ARYB2QO | -2 | 0.54296683 | 0.507632339 | 0.988135593 |
| AS2MVDL | -1 | 0.334167625 | 0.328039705 | 0.9421391 |
| ASFODQR | -2 | 0.568365176 | 0.536025995 | 0.975686733 |
| ATHBFWX | -2 | 0.51051866 | 0.464274324 | 0.972355348 |
| ATHKM6V | -2 | 0.164923601 | 0.083839621 | 0.940268849 |
| ATT9RR1 | -1 | 0.465272757 | 0.420144721 | 0.963062537 |
| AUCGUF3 | -2 | 0.562297842 | 0.490017124 | 0.946230275 |
| AUILKHG | -1 | 0.432987242 | 0.404546458 | 0.947574518 |
| AURCTAK | -1 | 0.482616562 | 0.468609646 | 0.979310345 |
| AUY8KYW | -1 | 0.646110521 | 0.590229301 | 0.99421391 |
| AV2GF3B | -1 | 0.218067321 | 0.206962728 | 0.972881356 |
| AW4EXXK | -1 | 0.407614929 | 0.380905892 | 0.948568089 |
| AWA2KJK | -2 | 0.504295305 | 0.469122208 | 0.960841613 |
| AWRBUQZ | -2 | 0.348714635 | 0.318765441 | 0.989246055 |

| | | | | |
|---|---|---|---|---|
| AX3KEW9 | -2 | 0.508648683 | 0.466224953 | 0.972998247 |
| AX6281V | -1 | 0.484778598 | 0.451470151 | 0.976914085 |
| AXCO7I9 | -2 | 0.405069604 | 0.354378135 | 0.989129164 |
| AXD3W8O | -1 | 0.550576436 | 0.535545964 | 0.969900643 |
| AXDWDEA | -1 | 0.205399804 | 0.16905398 | 0.976446523 |
| AXI1PBS | -2 | 0.572993673 | 0.52706524 | 0.979135009 |
| AY8TPMP | -1 | 0.687388919 | 0.665070586 | 0.992752776 |
| AYEFCWQ | -1 | 0.338694614 | 0.310862289 | 0.969842198 |
| AYVQUF1 | -2 | 0.501009881 | 0.452443735 | 0.973173583 |
| AYWIEKR | -2 | 0.475534987 | 0.442181758 | 0.983752192 |
| AZ2RYW7 | -2 | 0.462640402 | 0.419668547 | 0.990005845 |
| AZ35PI5 | -2 | 0.190317585 | 0.167486191 | 0.983284629 |
| AZIK4ZA | -2 | 0.627991625 | 0.571335667 | 0.982057276 |
| AZKZ0AI | -1 | 0.558288749 | 0.523446065 | 0.992694331 |

**Supplementary Table 2. Parameters in LagCI analysis for demo datasets.**

| program | time_tol (h) | step (h) | align_method | cor_method |
|---|---|---|---|---|
| wearable_analysis | 0.5 | 1/60 | linear | spearman |

| program | time_tol (h) | step (h) | align_method | cor_method |
| --- | --- | --- | --- | --- |
| omics_analysis | 12.25 | 30/60 | linear | spearman |
